# Effective repurposed antivirals against virus

**DOI:** 10.1101/2025.10.12.681938

**Authors:** Huiyuan Zhou, Sha Li, Lige Zhang, Li Liu, Tianyue Zhang, Yun Wang, Gaoyang Li, Huansheng Cao

## Abstract

The ongoing global monkeypox virus (MPXV) outbreak urgently needs effective medications, which be accelerated through drug repurposing. However, it is challenging to poinpoint protein targets. Here we introduce a novel method rooted in molecular evolutionary theory for quick drug target identification for MPXV. It identifies drug targets as positively selected genes of viral proteins which bind host proteins. From the identified targets, we select a top gene product OPG021 for virtural drug screening. One top-ranked drug nilotinib is experimentally shown to have a significant 69% of antiviral efficacy of the FDA-approved antiviral tecovirimat (TPOXX^®^ or ST-246). Higher binding affinity but not antiviral efficacy of repursposed drugs than FDA-approved drugs suggests the complexity of drug repurposing and underscores the importance of experimental validation. This innovative drug target identification strategy will contribute to combating the ongoing MPXV outbreak and other viral acute and chronic viral diseases.

## Introduction

Drug repurposing has great potential in quickly finding existing drugs to treat diseases, particularly infectious diseases ^1, 2^. This approach takes advantage of recently developed extensive drug databases ^2, 3, 4^ and advanced computational tools for virtual drug screening ^1, 5^. Despite these advancements, two critical challenges remain: the accurate and cost-effective identification of drug targets and untested efficacy of repurposed drugs. To date, drug target identification has been both inaccurate and expensive ^6^. Additionally, with only a limited number of candidate drugs having undergone experimental verification.

One common target-identifying approach involves leveraging viral-host protein-protein interactions (PPIs) ^6^. Although biologically sound, this bottom-up approach is both difficult and costly to execute. When performed experimentally ^7^ or computationally ^7, 8^, it often yields an overwhelming number of candidate targets. Another strategy involves identifying core genes among the genomes of various viral lineages ^9^, which are then screened based on specific biological properties, such as soluble, non-membrane-bound proteins. However, this approach also results in an excess number of candidate targets, complicating the selection process.

In this study, we modified existing drug identification methods by applying a simple criterion and applied it on MPXV, which has been the subject of a global outbreak (Moss, 2024; WHO, 2024) and declared a global health emergency by WHO on August 14, 2024 (doi: https://doi.org/10.1038/d41586-024-02607-y). So antiviruals against MPXV are desperately needed. Specifically, we began by identifying potential drug targets through viral-human PPIs and further screened the viral genes that have evolved under positive selection in MPXV which undergoes adaptive evolution ^10, 11^. Genes under positive selection in molecular evolution drive viral spread ^12^. These genes exhibit a signature frequency of nonsynonymous (amino acid-altering, *d*_*N*_) versus synonymous (silent, *d*_*S*_) substitutions between different lineages, with a *d*_*N*_/*d*_*S*_ ratio greater than 1, which can be computed using tools such as PAML^13^.

Using this approach, we identified and tested the gene OPG021, which encodes a host range factor, and discovered that the top-ranked drug nilotinib could potentially inhibit MPXV. *in vivo* tests on monkey cell line showed that they achieved a 72% and 100% inhibition rate against MPXV infection, relative to the FDA-approved antiviral medication tecovirimat.

## Results

### Viral-human protein-protein interaction

Following the design (Fig. 1), InterSPPI successfully identified 83% (152 out of 184) of the proteins encoded by the MPXV genomes, as detailed in Supplemental Table S1, with the ten most prominent proteins outlined in Table 1. Subsequent analysis revealed that the protein OPG021 (MPXVgp008) exhibits a nonsynonymous to synonymous mutation ratio (*d*_N_/*d*_S_) of 1.22, suggesting its coding gene is positively selected. Though its biological role is not thoroughly characterized, OPG021 is designated as a host range factor p28, soluble and not associated with membranes. Utilizing AlphaFold3, we inferred its tertiary structure (Fig. 2a), predicting its configuration to contain a KilA-N domain at the N-terminus and a RING-type zinc finger domain at the C-terminus.

**Table 1.** Top 10 gene products interacting with human proteins.

| Viral gene | Human gene | Binding score |
| --- | --- | --- |
| OPG021 | DHX9 | 0.304 |
| OPG200 | DYHC1 | 0.303 |
| OPG198 | DHX9 | 0.295 |
| OPG200 | UBR5 | 0.292 |
| OPG200 | SPTN1 | 0.289 |
| OPG022 | SPTN1 | 0.287 |
| OPG022 | DHX9 | 0.286 |
| OPG112 | ERBIN | 0.286 |
| OPG027 | ERBIN | 0.285 |
| OPG021 | DYHC1 | 0.284 |

**Fig. 1.**
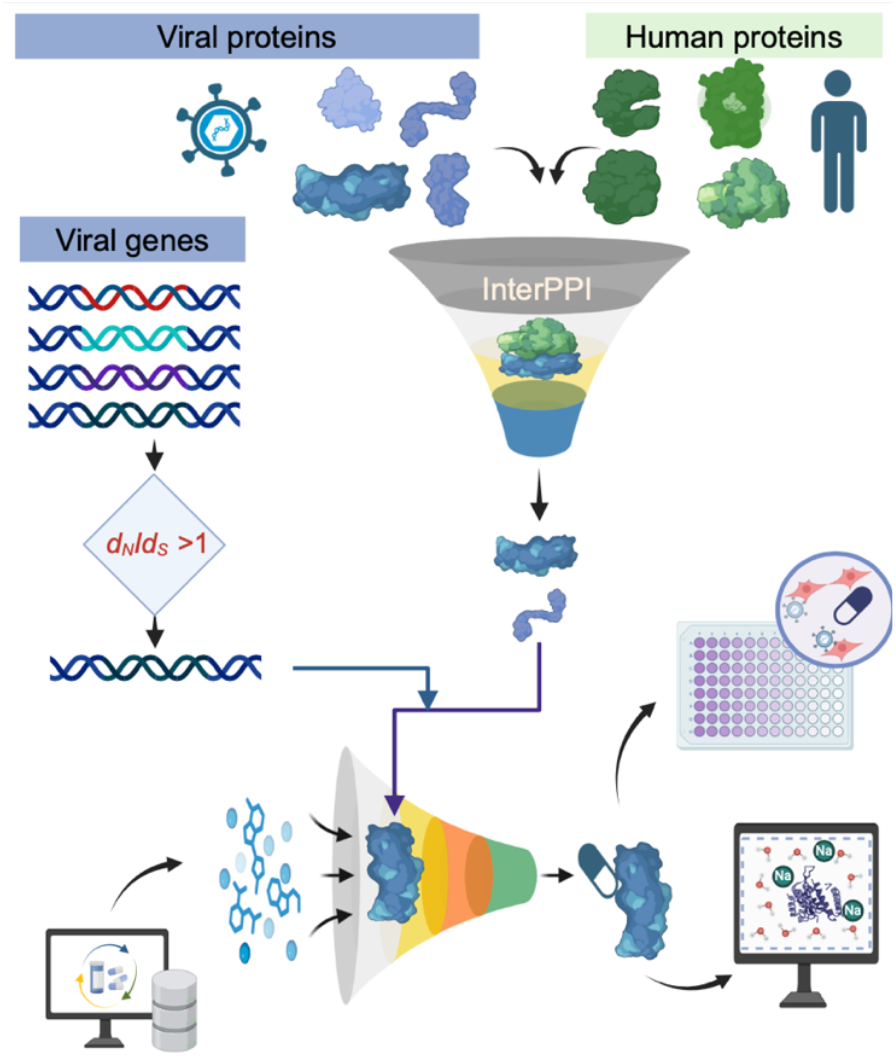
Overview of the complete drug repurposing process, including drug target identification and antiviral testing of repurposed drugs.

**Fig. 2.**
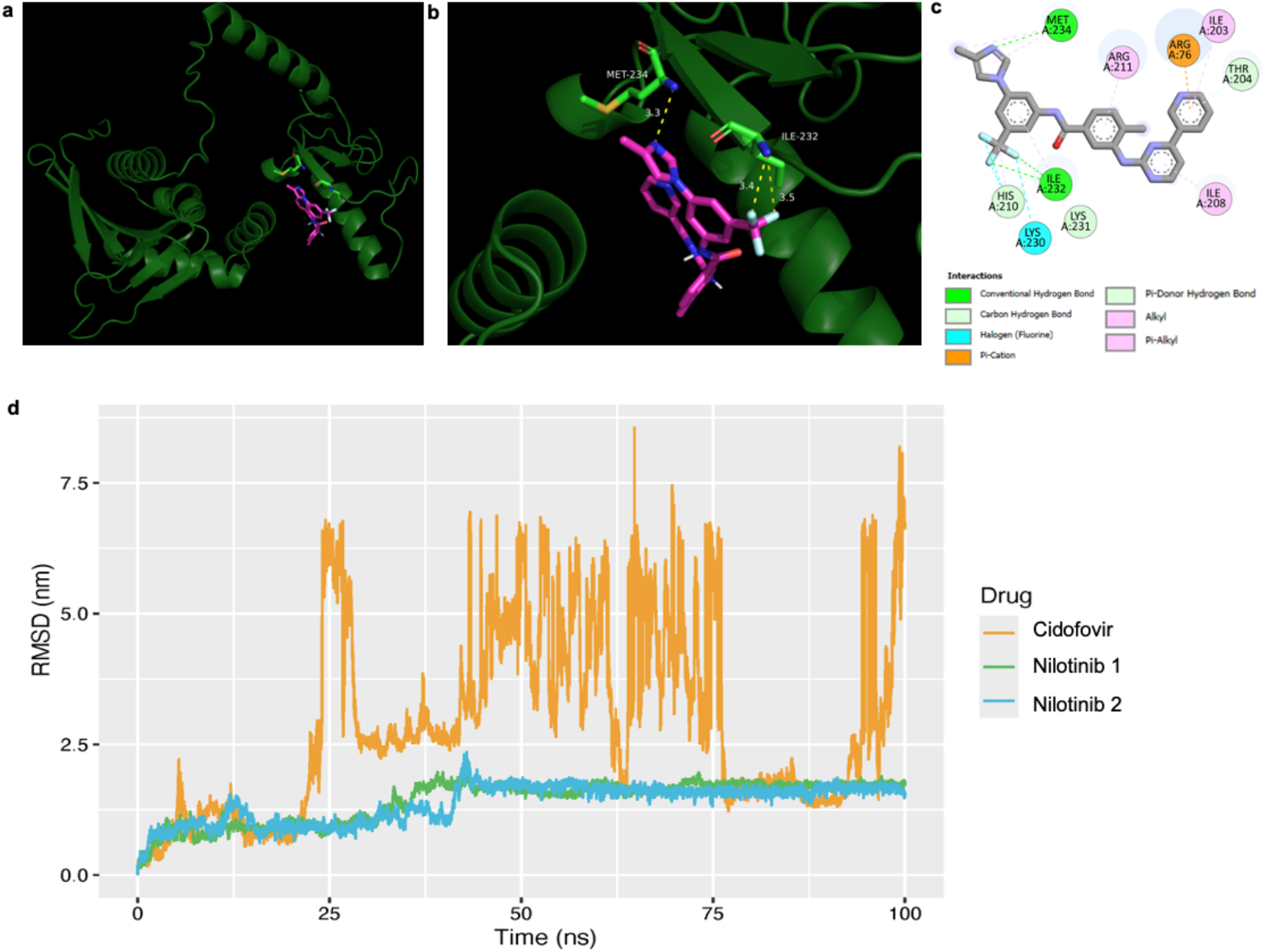
Drug-protein binding and molecular dynamics silumaiton. **a**: Global views of the docking sites for nilotinib on OPG021. **b**: Close-up views showing the hydrogen bonds between OPG021 and nilotinib. **c**: Specific bond types formed between OPG021 and nilotinib. **d**: Molecular dynamics simulation of the drug-protein complex over 100 ns in two runs, with cidofovir used as a negative control.

### Virtual drug screening

Through virtual drug screening, 345 potential inhibitors were identified as having stronger binding affinity than all three FDA-approved drugs (Table S2). Nilotinib was recognized the top candidates (Table 2), with the highest binding affinity of nilotinib (−8.91 kcal/mol), while the three FDA-approved drugs for MPXV treatment showed low binding affinity: tecovirimat (−7.42; ranked as 249), cidorvir (−5.03), and brincidofovir (−4.523).

**Table 2.** Top candidate drugs potentially inhibiting OPG021.

| EBC ID | Name | Binding affinity<br>(kcal/mol) |
| --- | --- | --- |
| EBC-11364 | Nilotinib | -8.91 |
| EBC-07301 | 2-(Naphthalen-1-yl)-1-(10H-phenothiazin-10-yl)ethan-1-one | -8.87 |
| EBC-191032 | Galicaftor | -8.78 |
| EBC-08915 | 1'-[(9-Ethyl-9H-carbazol-3-yl)methyl]-2,3-dihydro-1H-spiro[isoquinoline-4,4'-piperidin]-3-one | -8.67 |
| EBC-249047 | 3-[4-[(6,7-Dimethoxyquinazolin-4-yl)oxy]naphthalen-1-yl]-1-(4-[(morpholin-4-yl)methyl]-3-(trifluoromethyl)phenyl) urea | -8.56 |
| EBC-286055 | 2-[4-[11-Methyl-5-(pyridin-3-yl)-8-thia-4,6-diazatricyclo[7.4.0.0,2,7]trideca-1(9),2(7),3,5-tetraen-3-yl]piperazin-1-yl]-2-phenylacetamide | -8.53 |
| EBC-11409 | MLN8054 | -8.522 |
| EBC-08615 | Digoxin | -8.521 |
| EBC-148031 | N'-[(E)-(2-Hydroxynaphthalen-1-yl)methylidene]-3-(naphthalen-2-yl)-1H-pyrazole-5-carbohydrazide | -8.51 |
| EBC-13190 | NVP-BEZ235 | -8.496 |
| EBC-11433 | Lumacaftor | -8.49 |
| EBC-13180 | PHPS1 | -8.488 |
| EBC-303215 | 2-(3-[2-Oxa-6-azaspiro[3.3]heptane-6-carbonyl]benzenesulfonyl)-1,2,3,4-tetrahydroisoquinoline | -8.488 |
| EBC-157209 | URMC-099 | -8.449 |
| EBC-07988 | 1-[4-(3-Benzyl-5-phenyl-3H-[1,2,3]triazolo[4,5-d]pyrimidin-7-ylamino)phenyl]-3-(4-fluorophenyl)-urea | -8.44 |
| EBC-50104 | Mirk-in-1 | -8.428 |
| Control | Tecovirimat | -7.424 |
| Control | Cidofovir | -5.035 |
| Control | Brincidofovir | -4.523 |

### Binding affinity and molecular dynamics

Nilotinib appears to bind different sites on OPG021 (Fig. 2a). Close-up views showed these are two opposite sites and each has different hydrogen bonds to OPG021 (Fig. 2b). Nilotinib binds to one alpha helix at the site formed resides from ILE203 to MET234. A total of seven types of bonds are identified (Fig. 2c).

Further MD simulations was undertaken to assess a protein-drug complex’s conformational stability in its dynamic state during simulation, in terms of RMSD (root-mean-squared-deviation). Low differences in RMSD indicate low and consistent fluctuation between the ensembles and stable protein-ligand complexes. Among nilotinib and and three FDA-approved drug cidofovir, the top repurposed drug in two runs both achieved a relative equilibrium after 100 ns (Fig. 2d).

### In vivo antiviral test of repurposed drugs

Antiviral effects of nilotinib and hypericin were confirmed on monkey Vero E6 cell line in laboratory experiments. Using FDA-approved tecovirimat as a positive control (suppose its antiviral efficiency is 100%), we ran two batches of experiments (each with three replicates) and determined that nilotinib achieved 72% ± 25% and 93% ± 7% of antiviral efficacy. Cidofovir had antiviral efficiency of 70% ± 8% and 67 ± 5% in two batches of tests. We also determined the IC_50_ of nilotinib compared to Cidofovir and Brincidofovir, which was 31.91 µM, close to the IC_50_ for cidofovir 26.8 µM.

## Discussion

This study introduces a novel method for identifying drug targets against MPXV and validates its effectiveness through experimental results. The repurposed drug nilotinib demonstrates a high affinity for its target, OPG021, and exhibits significant antiviral effects, achieving a minimum efficacy of 72% and 100%. This outcome not only supports the new target-identifying method but also highlights a potential new treatment for the virus currently spreading.

The comparison between nilotinib and three existing antivirals reveals an intriguing aspect of drug repurposing: high ligand-protein binding affinity does not always translate to superior antiviral efficacy. Specifically, nilotinib has a binding affinity of −8.9 kcal/mol, higher than that of tecovirimat (−7.42 kcal/mol), brincidofovir (−5.03 kcal/mol), and cidofovir (−4.52 kcal/mol). Despite this, nilotinib’s inhibitory effect was only 72% of tecovirimat’s and brincidofovir’s, and comparable to cidofovir’s. Molecular docking simulations showed similar stability in the ligand-protein complexes among the four drugs, which reflects their binding affinities but not inhibitory effects. This discrepancy between the actual antiviral effect and *in silico* simulation underscores the importance of experimental validation in drug repurposing, reflecting its unexpected complexity ^14^.

This novel method for identifying drug targets bring an established molecular evolutionary perpective to the existing strength in protein-protein interaction ^15^. This positive selection has been observed in many viral outbreaks ^16^, even in the most recent SARS-COV-2 pandemics ^12^. Given the universal nature of this method in all infectious diseases, it can be applied all viral acute and chronic diseases, such as viral pneumonia ^17^.

Additionally, this study suggests that OPG021 might be a target for both FDA-approved tecovirimat and brincidofovir, as these drugs have similar chemical structures and antiviral effects. However, since their *in silico* binding to OPG021 is not as strong as that of nilotinib and hypericin, further experimental verification is required. Confirming this could also lead to improvements in these existing drugs.

In conclusion, the new method for identifying drug targets against MPXV proves effective, with nilotinib emerging as viable repurposed drugs. The study highlights the need for experimental validation in drug repurposing and suggests that OPG021 may be a target for tecovirimat and brincidofovir. Further research and testing will be essential to optimize these findings and combat the ongoing spread of MPXV, and other acute and chronic infectious diseases.

## Methods

### Viral-human protein-protein interaction

The viral proteins analyzed in this study were derived from MPXV Zaire-96-I-16 genome (accession: NC_003310.1), while the human genome data were sourced from the build GRCh38.p14 (NCBI RefSeq accession: GCF_000001405.40) ^18^. To predict interactions between viral and human proteins, the machine learning tool InterSPPI ^15^ was employed, utilizing default parameters for the analysis.

### Calculation of d_**N**_**/d**_**S**_

MPXV genomes were download from Nextstrain ^19^, which curated 736 representative genomes from NCBI (https://nextstrain.org/mpox/clade-IIb). The *d*_N_/*d*_S_ ratios for the genes within these genomes were calculated using the PAML software package ^20^.

### Virtual drug screening

The study utilized the Enamine Bioactive Collection library (https://enamine.net/compound-collections/bioactive-compounds), which contains 5,562 annotated compounds. This library was downloaded in March 2024. The 2D SDF files of the compounds were converted to 3D structures in MOL2 format using the RDKit Python toolkit (RDKit: Open-source cheminformatics. https://www.rdkit.org). For each ligand, 10 conformations were generated using the distance geometry method, corrected by the Experimental-Torsion Knowledge Distance Geometry (ETKDG). These conformers were then optimized with the MMFF94 force field, and the lowest energy conformer was selected and saved in MOL2 format. The MOL2 files were subsequently converted into PDBQT format using the Meeko package.

Since no crystallized structure of OPG021 was available, its protein structure was predicted using AlphaFold3 ^21^ Water molecules and non-relevant heteroatoms were removed, and polar hydrogens were added with Kollman charges assigned. The protein structure was then saved in PDBQT format, which is required for molecular docking with AutoDock.

Molecular docking was conducted using AutoDock Vina 1.2.5 ^22^. A blind docking approach was used, meaning the docking box covered the entire protein structure. For each protein-ligand interaction, 50 docking simulations were performed, and the exhaustiveness parameter was increased to 16 from the default value of 8 to enhance the search within the docking box (Trott & Olson, 2010). Ligands were ranked according to their binding energy across the 50 docking simulations. Compounds with binding energies lower than those of the control compounds were classified as hits and selected for further analysis. The controls used for comparison were three FDA-approved antivirals against MPXV: Tecovirimat, Brincidofovir, and Cidofovir.

### Simulation of molecular dynamics

Molecular dynamics (MD) simulations were conducted to evaluate the stability of protein complexes with nilotinib, tecovirimat, brincidofovir, and cidofovir. These simulations were carried out using GROMACS 2023.1 ^23^ and employed the CHARMM36 all-atom force field (July 2022 version), selected for its advanced optimization in protein-ligand interactions ^24^. Initial configurations for the simulations were based on the best-docked poses obtained from AutoDock Vina. Protein topology files were created using GROMACS’s built-in commands, while ligand topology files were generated via the CGenFF server (https://cgenff.com/). The protein-ligand complexes were embedded in dodecahedral unit cells and solvated with the SPC216 solvent model ^25^. The system was neutralized in charge by adding appropriate sodium or chloride ions. Following energy minimization to ensure system stability, convergence was confirmed before proceeding with further steps. The equilibration process involved sequential NVT (constant Number of particles, Volume, and Temperature at 300K) and NPT (constant Number of particles, Pressure, and Temperature at 300K) simulations, each lasting 100 ps. Production MD simulations were conducted for 100 ns, with 50,000,000 steps at 2 ps intervals for all ligands.

### In vitro antiviral effect test

The antiviral activity of nilotinib and hypericin against Monkeypox virus (MPXV) was assessed in Vero E6 cells, and compared with three controls: tecovirimat, brincidofovir, and cidofovir. Vero E6 cells were cultured at 37°C in a humidified incubator with 5% CO_2_, using Dulbecco’s Modified Eagle Medium (DMEM). All drug compounds were purchased from Sigma-Aldrich (St. Louis, MO, USA) and dissolved in dimethyl sulfoxide (DMSO) at concentrations of 10 mM and 20 mM, respectively.

The antiviral activity was assessed by the inhibition of the cytopathic effect (CPE) induced by MPXV. Vero E6 cells (10_4_ cells/well) were seeded in 96-well plates and treated with test drugs at a final concentration of 30 μM. Cells were subsequently infected with MPXV at a multiplicity of infection (MOI) of 0.1 plaque-forming units (PFU) per cell ^26^. Tecovirimat (ST-246) was used as a positive control, while untreated cell growth medium served as a negative control. After 72 hours, viral load was determined by quantitative PCR (qPCR) ^27^. Cells were harvested, and DNA was extracted using the MagaBio Plus Virus DNA/RNA Purification Kit III (BIOER, Hangzhou, China) according to the manufacturer’s instructions. Viral DNA was quantified by a TaqMan-based real-time PCR targeting the OPG087 gene region of the virus, using TaqMan Gene Expression Master Mix (Vazyme Biotech Co., Ltd., Nanjing, China). The following primers and probe were used: forward primer, 5′-CACACCGTCTCTTCCACAGA; reverse primer, 5′-GATACAGGTTAATTTCCACATCG; TaqMan probe, 5′-FAM-AAGCCGTAATCTATGTTGTCTATCGTGTCC-BHQ1. Antiviral efficacy was expressed relative to tecovirimat (positive control) as follows:

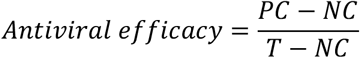

where *PC* and *NC* represent the positive and negative controls, and *T* represents the test drugs.

Next, EC_50_ values for each compound were determined by testing seven concentrations (100, 32, 10, 3.2, 1, 0.32, and 0.1 μM). All other procedures, including qPCR-based CPE analysis, followed the protocol from the antiviral assays. Dose-response curves were generated using a four-parameter logistic model (variable-slope, nonlinear regression), and EC_50_ values were calculated using Prism 9.0 (GraphPad Software, MA, USA).

## Acknowledgements

The work was supported by Kunshan Municipal Research Grant.

## Author contributions

L.Z., L.L., and T.Z. performed InterSPPI and *d*_N_/*d*_S_ analyses, HZ performed drug virtual screening and molecular dynamics simulation, S.L. performed drug testing, H.C. and G.L. conceived the project. G.L., H.C. and Y.W. designed experiments and wrote manuscript. All authors reviewed final manuscript.

## Code availability

The code for this study can be found at https://github.com/lgyzngc/mpxvdrug.

